# Looking into working memory to verify potential targets during search

**DOI:** 10.1101/2024.10.28.620583

**Authors:** Sisi Wang, Freek van Ede

## Abstract

Finding what you are looking for is a ubiquitous task in everyday life that relies on a two-way comparison between what is currently viewed and internal search goals held in memory. Yet, despite a wealth of studies tracking visual verification behavior among the external contents of perception, complementary processes associated with visual verification among internal contents of memory remain elusive. Building on a recently established gaze marker of internal visual focusing in working memory, we tracked the internal-inspection process associated with confirming or dismissing potential targets during search. We show how we look back into memory when faced with external stimuli that are perceived as potential targets and link such internal inspection to the time required for visual verification. A direct comparison between visual verification among the contents of working memory or perception further revealed how verification in both domains engages frontal theta activity in scalp EEG, but also how mnemonic verification is slower to deploy than perceptual verification. This establishes internal verification behavior as an integral component of visual search, and provides new ways to look into this underexplored component of human search behavior.

## Introduction

A ubiquitous task in everyday life is to find what you are looking for, a process commonly referred to as visual search. Models of visual search naturally invoke the notion of a comparison of what is currently viewed to a search target or “template” held in mind, in visual working memory ^1–9^. This comparison process allows to verify whether something externally in view does or does not match one’s internal search goal held online in memory.

Visual verification behavior is classically characterized by how we explore and inspect external visual stimuli: we gravitate toward and further inspect external visual stimuli that resemble what we are looking for ^1,10–17^. Critically, because visual verification is fundamentally about finding a *match* between internal *and* external visual information, the verification process is likely to involve a two-way stream: just like internal memory templates may invite and prompt further inspection of resembling external stimuli, so too may external stimuli invite and prompt further inspection of the internal search-target representations that they resemble, in service of target verification (i.e., to confirm or dismiss a match). Yet, evidence directly tracking such internal inspection processes remain scarce and key questions remain unaddressed.

Consistent with the notion of an internal inspection serving visual verification, at least two prior studies have shown that when an external visual target has to be compared to multiple visual objects (templates) held in working memory, we return to the memorized location associated with the memory template ^18–20^. While these studies are consistent with the sketched internal-verification process, these studies have left two foundational questions unaddressed. First, if this process reflect internal inspection in service of visual verification, it should not only occur when a visual target matches a mnemonic search goal (as these studies showed), but also when it merely *resembles* what you are looking for (i.e., a potential target). Because aforementioned studies only showed the former, it has remained ambiguous whether the reported findings reflected the verification process itself (going back in mind *in order to* verify) or the *outcome* of this process (going back in mind *after* a target has been verified). Second, this putative internal-inspection-for-verification process has yet to be linked to actual verification behavior. If mnemonic inspection is relevant to visual verification, it should vary as a function of the time needed to verify whether or not a visual stimulus matches a search target in working memory.

To address these foundational open questions, we developed a dedicated task in which participants judged whether centrally presented visual stimuli were either identical to (target) or merely resembled (non-target lures) one of two visual representations held in working memory. By using spatially separated memory targets, we could isolate attentional allocation associated with internal visual inspection in working memory (following the same logic as in ^18–24^). We here tracked such internal inspection within the spatial layout of visual working memory through spatial biases in microsaccades that we have recently uncovered as a powerful signal to track internal visual focusing ^23–30^. Critically, by also including lures that resembled either memory object, we could track mnemonic verification behavior not only following actual visual targets (as in ^18,19^) but also following non-target lures that were perceived as potential targets and thus invited further mnemonic inspection. This uniquely revealed how internal visual verification among the contents of working memory occurs not only when a visual target matches a mnemonic search goal (as in ^18–20)^, but also when it is perceived as a potential target because it resembles a search goal – confirming the interpretation of an internal inspection-for-verification process. Moreover, we here for the first time link this internal inspection process to observed visual verification behavior, showing it is particularly pronounced when the verification process takes more time to resolve.

In addition, we unveil important commonalities as well as differences when engaging visual verification by further inspecting the contents of perception (as widely studied previously) or working memory. We show how perceptual- and mnemonic-verification processes similarly engage frontal theta activity in scalp EEG measurements (cf. ^31–38^), but also how it takes more time to deploy mnemonic verification among the contents of working memory as opposed to the more widely studied perceptual verification among the contents of perception.

## Results

Human observers performed two version of a visual-verification task that required verifying whether a visual stimulus on the screen matched either of two internal visual representations held in working memory (“mnemonic verification”; **Fig. 1a**), or whether a single visual representation held in working memory matched either of two external visual stimuli on the screen (“perceptual verification”; **Fig. 3a**).

**Figure 1.**
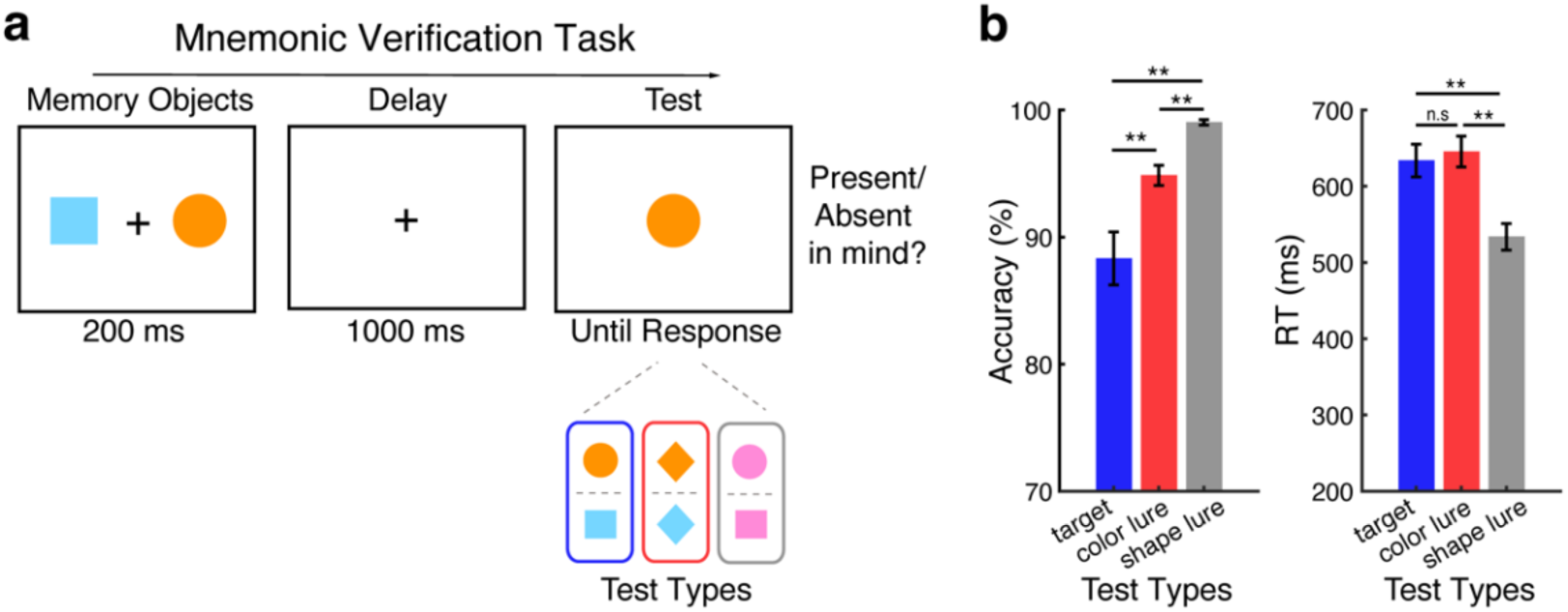
Visual target verification depends on perceived similarity between memorized and perceived objects. **a**) Top panel: schematic of the mnemonic-verification task. Participants memorized two visual objects (encoded to the left and right of fixation), and reported whether a subsequent centrally presented test stimulus was identical to either memory object (i.e. present/absent). Bottom panel: possible test-stimulus types. Test stimuli could be actual targets (color and shape match with one of the memory objects, as illustrated in the blue frame), color lures (only color match with one of the memory objects, as illustrated in the red frame) or shape lures (only shape match with one of the memory objects, as illustrated in the gray frame). Color and shape lures required an ‘absent’ report. Targets and lures only matches one or both features with one of the memory objects (i.e., the left or the right object), and never with the other. **b**) Response accuracy (left panel) and response time (right panel) across test types. Error bars represent ±1 SEM. ** represents *p* < 0.001, n.s represents not significant.

Our primary focus was on tracking whether mnemonic verification engages re-focusing (inspecting) specific mental representations for verification and to assess whether this occurs even when these representations are not actual targets, but are perceived as *potential* targets. To this end, we defined targets by color-shape pairs, but included “lures” (non-targets) that matched one of the memory objects only in color or only in shape (**Fig. 1a**; red and grey Test Types). Our rationale was the following: because lures *look like* potential targets, they may trigger internal inspection of similarly looking working-memory content to verify or dismiss a potential match.

To study such internal “in-mind” visual verification, we presented memory objects to the left and right while presenting the external comparison stimulus centrally (**Fig. 1a**). This enabled us to leverage spatial biases in fixational eye movements (microsaccades) to isolate internal attentional deployment within the spatial layout of working memory (as in ^23–30)^. Our complementary perceptual-verification task (**Fig. 3a**) – which is more established ^10–12,14–16,18,19,21^ – served mainly as a reference for interpreting our key mnemonic-verification findings. We note how both tasks engage matching mnemonic with perceptual information, but refer to these tasks as tracking “mnemonic” and “perceptual” verification respectively by virtue of our ability to experimentally isolate and track further visual inspection in working memory (mnemonic verification) or in perception (perceptual verification), as uniquely tagged by spatial location.

In what follows, we first provide eye-movement evidence for mnemonic (in-mind) verification that occurs not only for actual targets, but also for lures that appear like potential targets. We show how such internal inspection occurs particularly when the verification is non-trivial and takes time. We next delineate similarities and differences between this uncovered mnemonic verification behavior and the verification of potential perceptual targets, which is more established. Finally, we report EEG evidence for a common neural process mediating visual verification among the contents of working memory and among the contents of perception.

### Visual verification depends on perceived similarity between memorized and visible objects

Figure 1b shows behavioral performance on our mnemonic-verification task. While participants found it easy to dismiss shape lures (that always had a distinct color from the two memory objects) as non-targets, participants found it more difficult to reject color lures, or to confirm actual targets. This was confirmed statistically. Repeated measures ANOVAs revealed a significant main effect of Test Type (target vs. color lure vs. shape lure) for both accuracy (*F*(2,48) = 21.317, *p* < 0.001, η² = 0.470) and RT (*F*(2,48) = 78.202, *p* < 0.001, η² = 0.765). Post-hoc paired sample t-tests confirmed higher accuracy when the tested objects were shape lures, compared to color lures or actual targets (*t*(24) = 5.875, *p* < 0.001; *t*(24) = 5.351, *p* < 0.001, respectively), but also showed higher accuracy following color lures than following actual targets (*t*(24) = 3.402, *p* = 0.007). For RT, we also confirmed that people found it easier to reject shape lures compared to color lures and actual targets (*t*(24) = -12.167, *p* < 0.001, *t*(24) = -8.604, *p* < 0.001, respectively), while color lures and actual targets did not differ (*t*(24) = 1.419, *p* = 0.506).

While the observed lower accuracy for actual targets than for color lures may at least partially be explained by a response bias toward the ’different’ response option (given two-third of trials contained lures and only one-third targets), this cannot explain the observed differences between color and shape lures, as these lure-types were always equally likely.

Together, these data show that while shape lures were easy to dismiss, any visual stimulus that matched the color of either visual representation in mind – an actual target or a color lure – was more difficult to arbitrate and engaged a process that took additional time, as evidenced by RT. As we show next, such color-matching stimuli called upon internal inspection of the color-matching memory object in service of arbitrating color lures from actual targets.

### Spatial saccade bias tracks internal inspection for potential visual matches

To track the inspection of internal visual memory objects in service of visual verification, we capitalized on our recent findings that have uncovered how attentional allocation within the spatial layout of visual working memory can be “read out” from spatial biases in fixational gaze behavior ^23–30^, even in the presence of a large centrally presented stimulus ^24,26,29^ as was also the case here. By presenting our memory objects to the left and right, but our test stimulus centrally (Fig. 1a), we could use such spatial biases to isolate internal attentional focusing, as a marker of mnemonic inspection associated with the internal verification process.

Consistent with our behavioral results, we observed robust spatial saccade biases (Fig. 2) reflecting internal inspection following both targets and color lures, but not following shape lures that were less likely to be perceived (confused) as potential targets. Specifically, we found more saccades in the direction toward vs. away from the location of the visual memory representation that matched the target (Fig. 2a, left; cluster *p* = 0.011), or whose color matched the color lure that was perceived as a potential target (Fig. 2a, middle; cluster *p* = 0.034). We present an overlay of the spatial saccade biases across our three conditions in **Supplementary Figure S1a**.

**Figure 2.**
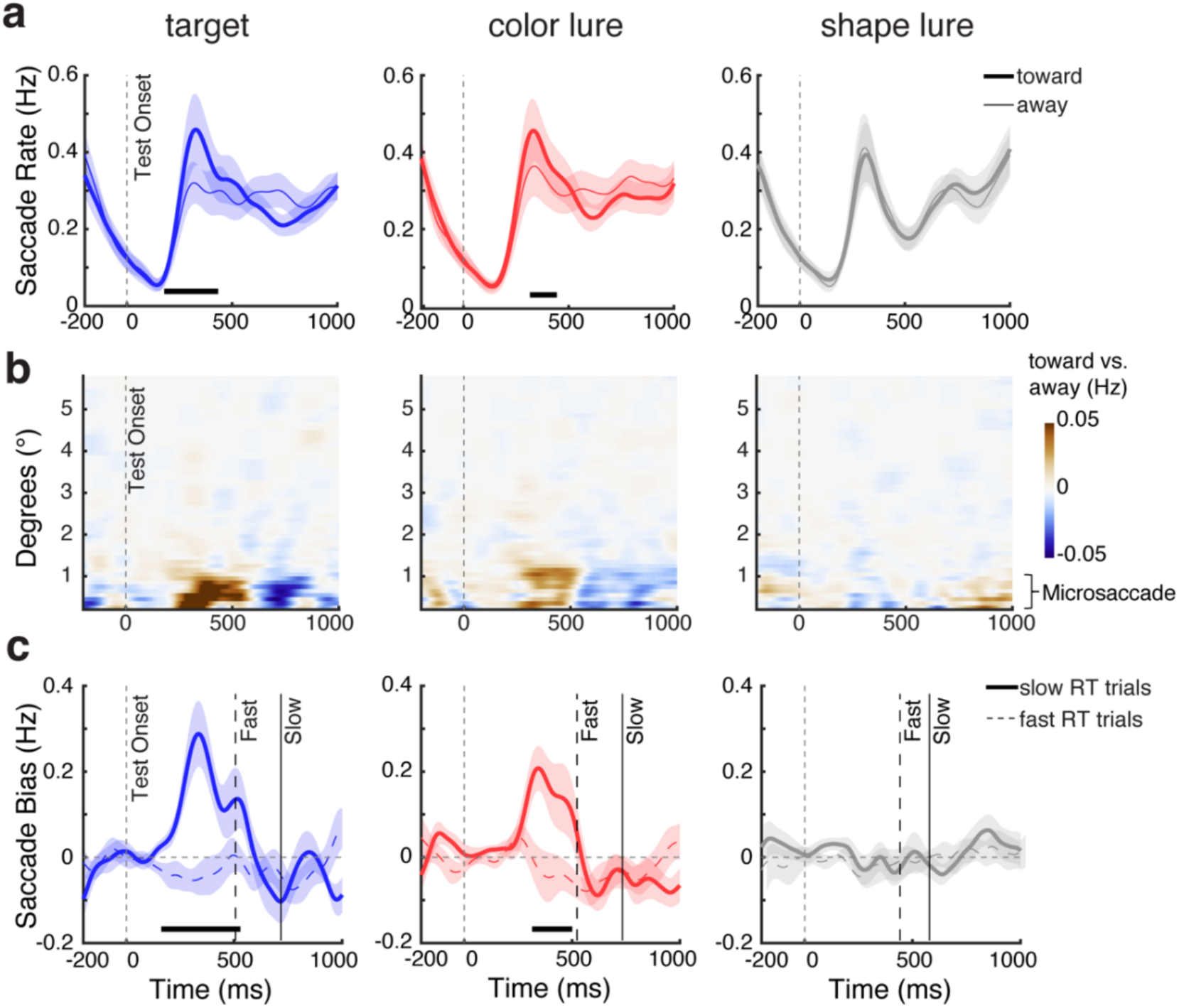
Spatial saccade bias tracks internal inspection following potential visual matches and is particularly pronounced when verification takes time. **a)** Saccade rates (in Hz) separately for toward (bold lines) and away (thin lines) saccades relative to the original (encoded) location of the memory object that matched the central test stimulus, when the test was the actual target (left), color lure (middle), or shape lure (right). **b)** Spatial saccade bias (the diTerence between toward and away saccades, with red colors denoting more toward saccades) as a function of saccade size (y axis) and time (x axis) across test conditions. This shows how the spatial saccade bias was predominantly driven by saccades below 1 visual degree, consistent with a bias in microsaccades. **c)** Spatial saccade bias (toward minus away) in trials with slow verification (RT > median RT; solid bold lines) and fast verification (RT < median RT; dashed thin lines) across test conditions. The solid and dashed vertical lines indicate the median RTs (averaged across participants) in the depicted conditions. The thick black horizontal lines indicate significant diTerence clusters. Shadings represent ± 1 SEM across participants (n = 25).

Consistent with our prior studies on this gaze marker of internal focusing ^24,26,27^, this saccade bias – quantified as the difference in rate between toward vs. away saccades – was driven predominantly by small saccades within the classical microsaccades range (Fig. 2b). Moreover, this saccade bias was observed prior to reporting (see Fig. 2c), consistent with an internal-inspection process serving the ensuing report decision.

While we found clear evidence for internal attention shifts triggered by targets and color lures that were perceived as potential targets – thus prompting further inspection – we found no evidence for similar effects following shape lures (Fig. 2, right column). This likely reflects the fact that shape lures were easy to dismiss based on their unique color, without requiring further inspection. This is consistent with our behavioral data showing that participants rarely confused shape lures and were fast to dismiss them as non-targets.

### Internal inspection is particularly pronounced in slow verification trials

We next reasoned that if the identified gaze bias reflects internal inspection in service of mnemonic verification, it should be particularly prominent in those trials where the additional verification is required – but may come at the expense of additional *time* required for verification. In other words, it may be particularly pronounced in slow verification trials. To address this prediction, we separated trials with fast and slow verification decisions in our task, by performing a median split on RT within each condition. As shown in Figure 2c, this confirmed that the spatial gaze bias – as our marker of internal inspection – was pronounced particularly in those trials in which it took participants more time to verify whether the external stimulus matched the relevant internal representation or not. This was the case both when the stimulus was the actual target (Fig. 2c, left; slow vs. fast cluster *p* < 0.001), or a colour lure that was perceived as a potential target (Fig. 2c, middle; slow vs. fast cluster *p* = 0.005).

We replicated these key eye-tracking findings in a second experiment that contained two additional conditions (as described in our **Supplementary Material** and visualized in **Fig. S2**).

### Mnemonic verification is similar but also distinct from perceptual verification

To relate our mnemonic verification findings to the broader existing literature, we also examined behavioral and gaze patterns during a complementary perceptual-verification task (Fig. 3a) of which similar tasks have been used in ample prior research (e.g., ^10–12,17–19,22,39^). While this task again required finding a match between internal and external visual stimuli, this time we used one central memory object and two potential perceptual targets that were now presented to the left and right. Accordingly, in this set-up, spatial gaze biases signal perceptual inspection among the sensory contents of perception (rather than mnemonic inspection among the memorised contents of working memory as in the preceding results).

**Figure 3.**
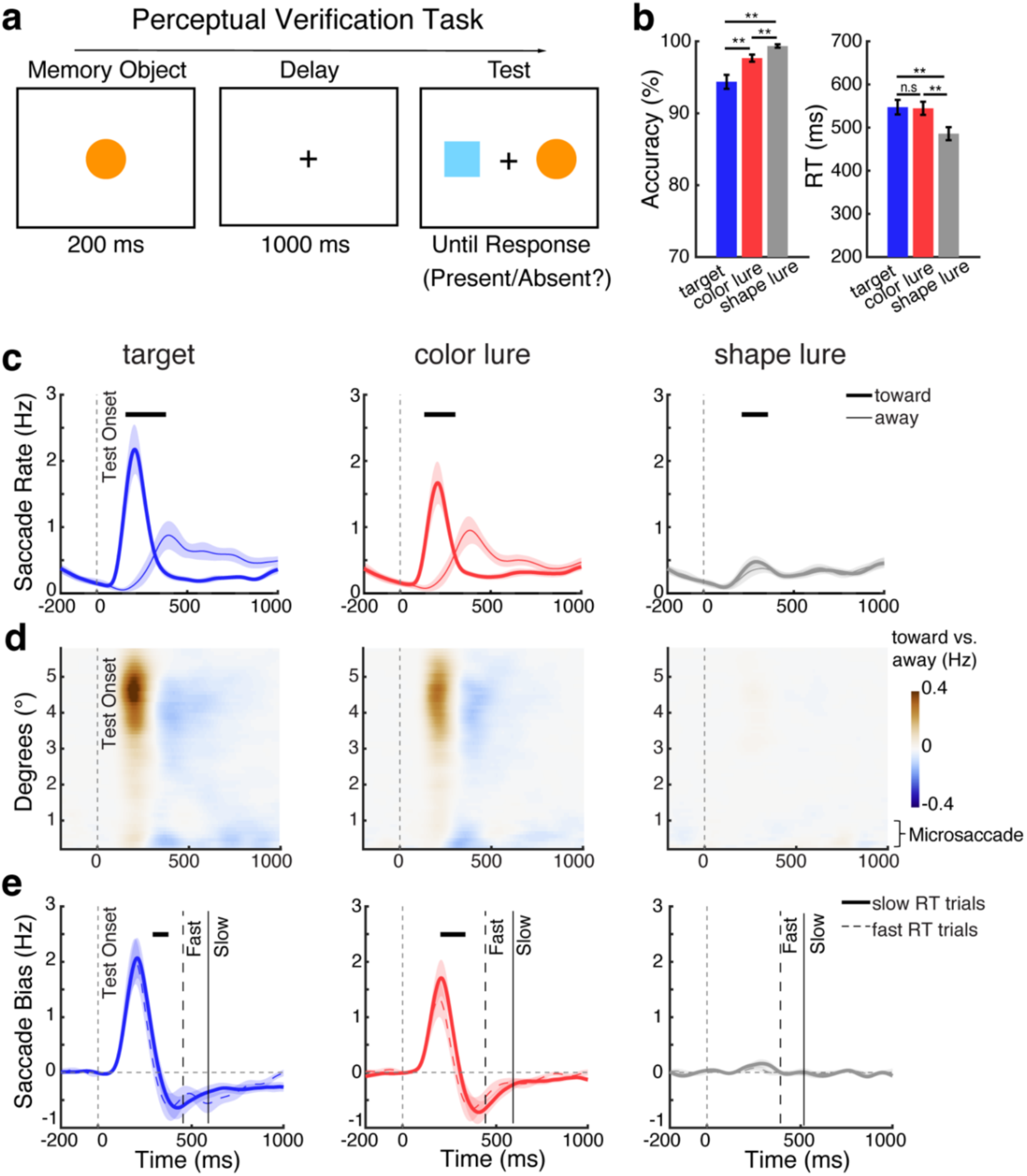
Task, behavior, and gaze in the perceptual-verification task. **a)** Schematic of the perceptual-verification task. Participants memorized one centrally presented visual object, and reported whether it was present or absent in a subsequent test array containing one left and one right visual stimulus. **b)** Response accuracy (left panel) and response time (right panel) across test types. Error bars represent ±1 SEM; ** represent *p* < 0.001; n.s represents not significant. c) Saccade rates (in Hz) separately for toward (bold lines) and away (thin lines) saccades relative to the location of the test stimulus that matched the memorized object, when one of the test stimuli was the actual target (left), color lure (middle), or shape lure (right). d) Spatial saccade bias (the diTerence between toward and away saccades, with red colors denoting more toward saccades) as a function of saccade size (y axis) and time (x axis) across test conditions. In this version of the task, the spatial saccade bias was driven by larger macro saccades associated with looking at the matching external visual stimulus that was centred at 5.4 degrees. e) Spatial saccade bias (toward minus away) in trials with slow RT (RT > median RT; solid bold lines) and fast RT (RT < median RT; dashed thin lines) across test conditions. The solid and dashed vertical lines represent the median RTs (averaged across participants) in the depicted conditions. The thick black horizontal lines indicate significant diTerence clusters. Shading represents ± 1 SEM across participants (n = 25).

Similar to target verification in working memory, we again found that shape lures were easiest to dismiss, compared to color lures and actual targets (Fig. 3b). This was confirmed statistically. Repeated measures ANOVAs revealed significant main effect of Test Type for both accuracy (*F*(2,48) = 27.843, *p* < 0.001, η² = 0.537) and RT (*F*(2,48) = 43.447, *p* < 0.001, η² = 0.644). Post-hoc paired sample t-tests confirmed higher accuracy when the tested objects were shape lures, compared to color lures or actual targets (*t*(24) = 4.218, *p* = 0.001; *t*(24) = 5.588, *p* < 0.001, respectively), but also showed higher accuracy following color lures than following actual targets (*t*(24) = 5.032, *p* = 0.007). For RT, we also confirmed that participants found it easier to reject shape lures compared to color lures and actual targets (*t*(24) = -13.219, *p* < 0.001; *t*(24) = -6.722, *p* < 0.001, respectively), while color lures and actual targets did not differ (*t*(24) = -0.338, *p* = 1.000).

When considering gaze, we also replicated clear spatial saccade biases to targets as well as to lures – whose locations were now not in working memory but on the screen (Fig. 3c). While this was again clearest for actual targets (cluster *p* = 0.006) and color lures (cluster *p* = 0.005), the bias also reached significance for shape lures (cluster *p* = 0.024). An overlay of the spatial saccade biases between conditions can be found in **Supplementary Figure S1b.** Unlike during mnemonic verification, where we found a spatial bias in the microsaccade range, during perceptual verification this bias was driven by larger (regular) “macro” saccades (Fig. 3d), consistent with the fact that now there were external visual stimuli (rather than internal memory representations) to inspect. We further replicated a relation between the magnitude of this bias and participant’s RT (Fig. 3e) following actual targets (Fig. 3e, left; slow vs. fast cluster *p* = 0.004) and color lures (Fig. 3e, middle; slow vs. fast cluster *p* < 0.001), again showing a larger bias in slow verification trials. However, unlike what we observed for mnemonic inspection, perceptual inspection was pronounced even when participants were fast.

To further elucidate the differences between visual target verification by further inspecting working memory or perception, we directly compared the gaze patterns between the mnemonic and perceptual tasks versions. Comparing the normalized saccade bias (peak normalized to be in the same range), we found that the saccade bias started substantially later during mnemonic inspection among the contents of working memory compared to during perceptual inspection among the contents of perception, following both actual targets and color lures (Fig. 4a**,b**). This was confirmed statistically using a jackknife test on the onset latency (quantified as the first value reaching 50% of the peak) both following actual targets (Fig. 4a; 261.6 ms for mnemonic inspection vs. 144.5 ms for perceptual inspection, *p* < 0.001) and following color lures (Fig. 4b; color lure: 250.2 ms for mnemonic inspection vs. 142.0 ms for perceptual inspection, *p* = 0.005).

**Figure 4.**
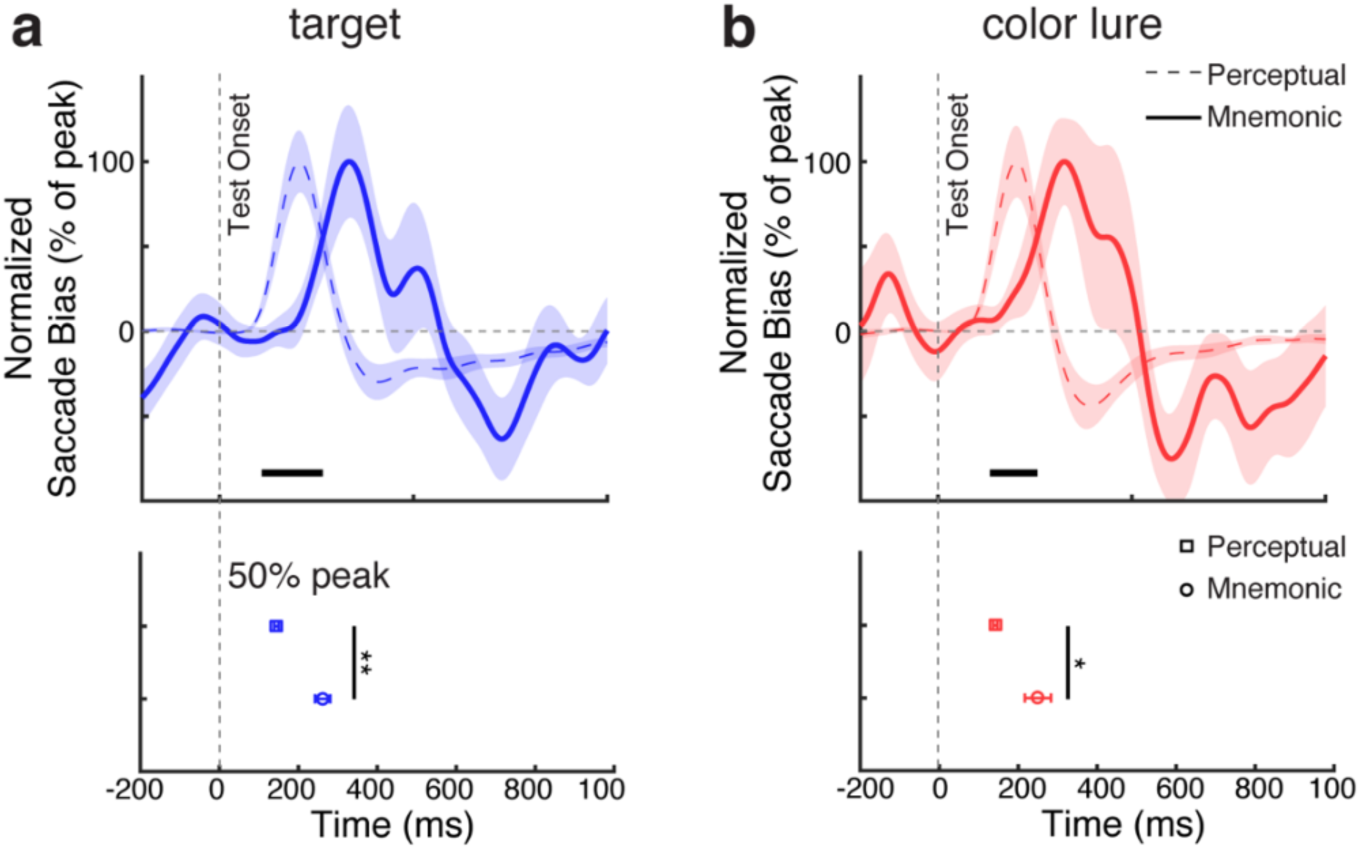
Mnemonic inspection is initiated slower than perceptual inspection in service of visual verification. Top panels: Normalized spatial saccade biases (toward vs. away) associated with mnemonic inspection (solid bold lines) and perceptual inspection (dashed thin lines) following onset of a) actual targets and b) color lures. The thick black horizontal lines indicate significant diTerence clusters. Shading and error bars represent ± 1SEM across participants (n = 25). To facilitate comparison between mnemonic and perceptual verification processes, saccade-bias time courses were normalized to their peak value. Bottom panels: Onset latency of the spatial gaze bias, defined as the first sample reaching 50% of the peak, during perceptual (squares) and mnemonic (circles)verification. **, and * represent *p* < 0.01, and < 0.05, respectively.

### Frontal midline theta activity track both mnemonic and perceptual verification

Our behavioral and gaze data suggested robust visual inspection processes following actual targets and color lures, but not following shape lures. Presumably, in our task, shape lures could be dismissed without further inspection, purely based on their distinctive color. Here, we leveraged this observation to investigate the neural dynamic of the inspection-for-verification process using Electroencephalography (EEG). The relative lack of a putative inspection-for-verification process in the shape-lure condition enabled us to use this condition as a reference condition to contrast to the other two conditions that did show clear evidence of such verification processes.

This revealed a relative increase in frontal midline theta (∼3-7 Hz) activity in the two conditions that showed the visual-inspection-for-verification process of interest. This was the case during both mnemonic verification (Fig. 5a; cluster *p* = 0.035 and 0.003 for the target and color-lure conditions, respectively) and perceptual verification (Fig. 5b; cluster *p* = 0.006 and 0.005 for the target and color-lure conditions, respectively). This frontal theta increase happened before response onset (see median condition-specific RTs indicated by the vertical dashed lines in Fig. 5a**,b**), consistent with reflecting the verification process preceding the report.

**Figure 5.**
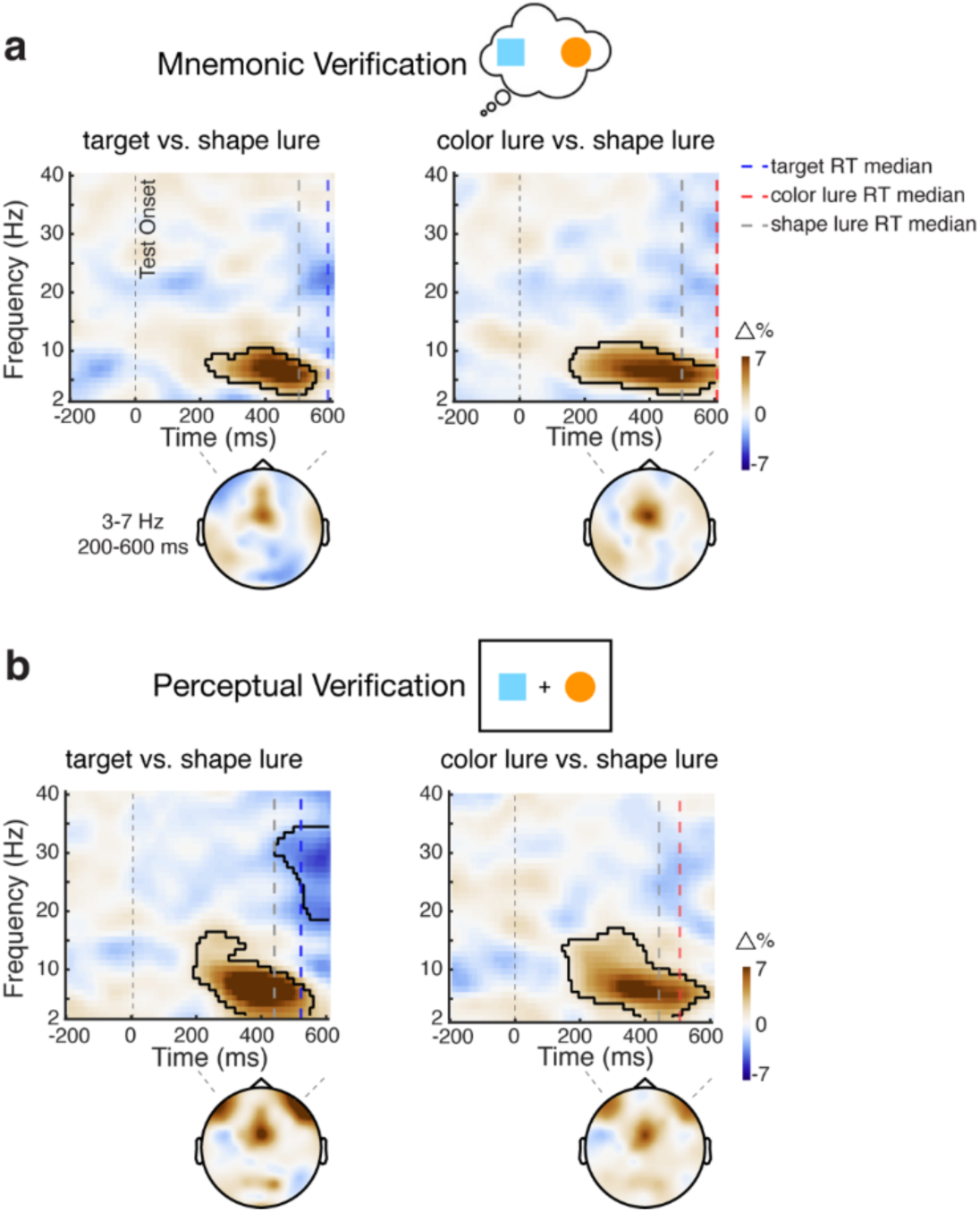
Frontal midline theta activity tracks visual verification among the contents of working memory and perception . **a**) Mnemonic verification: time-frequency maps of the percentage change in spectral power following targets (left panels) and color lures (right panels) in comparison to shape lures that here served as the neutral reference condition. The colored vertical dashed lines represent median RT for each condition (averaged across participants). Blue, red, and gray lines represent median RT for target, color lure and shape lure, respectively. Black outlines indicate significant clusters. Topographies show the diTerence in power averaged over 3-7 Hz and 200-600 ms after the test onset. **b)** The same conventions as in panel a, but for the data from the perceptual-verification task.

Additionally, our data in Figures 2c and **3e** suggested that this verification process (in the target and color-lure conditions) was particularly pronounced in trials with slow responses where, presumably, the verification process was more demanding and thus time consuming. Accordingly, if frontal theta activity supports the verification process, it should also be more pronounced in slow trials. This is precisely what we found, with stronger frontal theta activity in slow-RT compared to fast-RT trials (Fig. 6a, mnemonic verification: cluster *p* = 0.024 and 0.028 for the target and color-lure conditions, respectively; Fig. 6b, perceptual verification: cluster *p* < 0.001 and 0.004 for the target and color-lure conditions, respectively).

**Figure 6.**
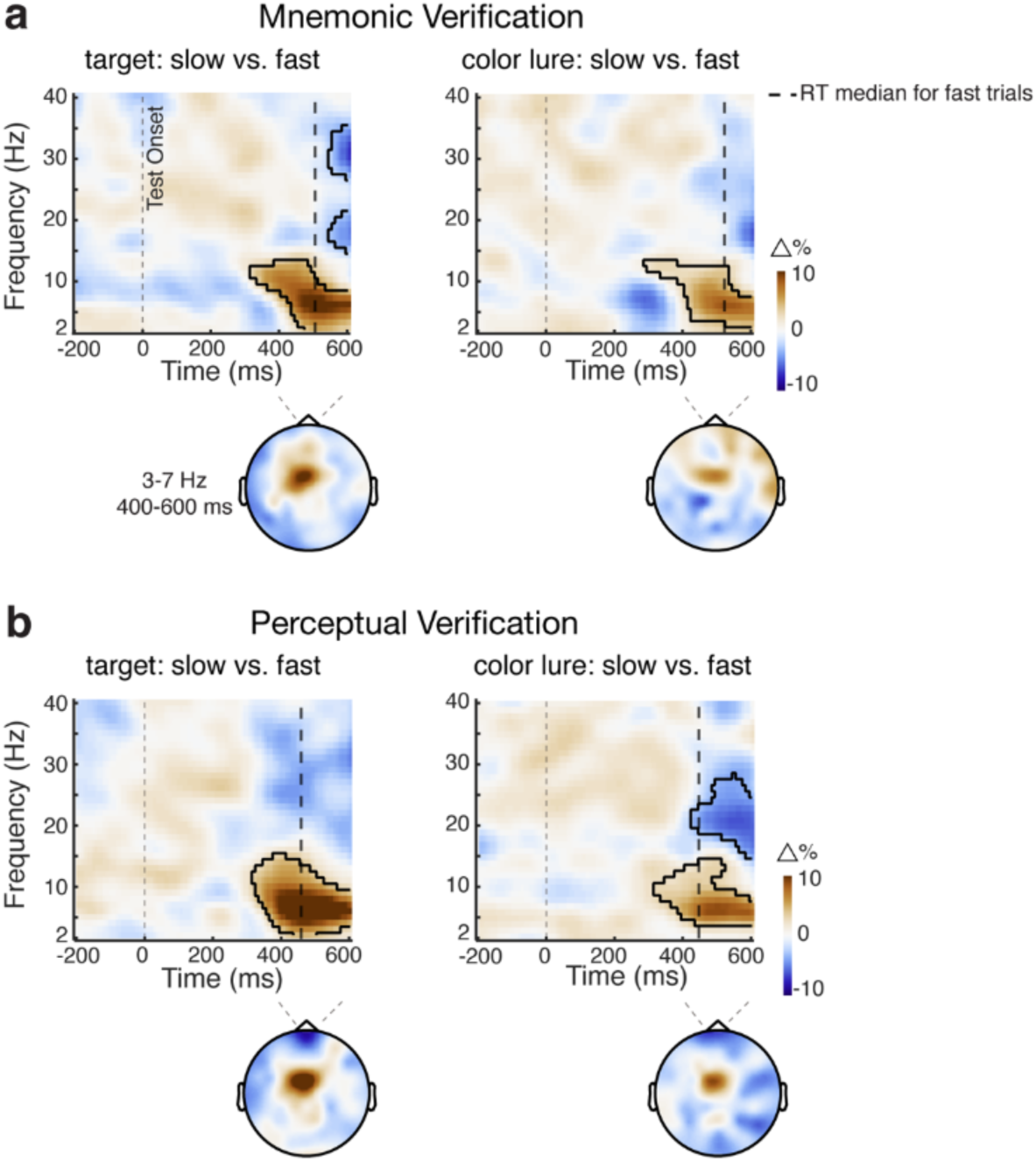
Frontal midline theta activity is particularly pronounced when verification takes time. **a**) Mnemonic verification: time-frequency maps of the percentage change in spectral power between slow minus fast verification trials (separated by a median split), within the target (left panels) and color lure (right panels) verification conditions of interest. The black vertical dashed lines represent median RT for fast verification trials for each condition (averaged across participants). Black outlines indicate significant clusters. Topographies show the diTerence in power averaged over 3-7 Hz and 400-600 ms after the test onset. **b)** The same conventions as in panel a, but for the data from the perceptual verification task.

## Discussion

Models of visual search – finding what we are looking for – naturally invoke the notion of a comparison between internal memory representations (mnemonic search goals) and incoming external sensory information ^3,7,8,40^. Accordingly, verification of potential matches (to confirm or dismiss a visual target as a match) may not only call upon further inspection among the external contents of perception (as e.g. in ^10,11,17,41^), but also among the internal contents of working memory. Yet, studies directly tracking the latter have remained scarce and have left key questions unaddressed. By studying human gaze behavior we provide evidence that humans look back into memory during visual search. Consistent with a verification process, we show that such mnemonic inspection occurs not only when a visual target matches a mnemonic search goal but also when it resembles a *potential* search goal that prompts further internal inspection. We further show how this process scales with observed verification behavior, being particularly pronounced when it takes more time to resolve a potential match. These findings underscore the relevance of internal verification processes in visual search and provide new ways to expose and track this underexplored component of visual search.

At least two prior event-related potentials (ERPs) studies have reported how visual search – finding a match between visual information in mind and world – can engage shifts of attention to matching internal representations held within the spatial layout of visual working memory ^18,19^ (for a complementary behavioral demonstration, see also ^20^). Our findings build on and extend this work in three fundamental ways. First, we show that such mnemonic inspection serving verification also occurs for *potential* targets that merely resemble what we are looking for. Because aforementioned studies could only track internal inspection following match-targets, it had been left unaddressed whether the reported findings reflected the verification process itself (inspection serving verification), or the consequence of having verified the target. Our study now provides clear evidence for the former interpretation. Second, we for the first time relate this internal-inspection process to observed verification behavior, showing it is particularly pronounced when the verification process requires more time to resolve. Finally, our data showcase the usefulness of gaze as a powerful complementary measure to track such internal attentional focusing during search – that we here interpret as a mnemonic inspection process serving verification. Such “looking into memory” may benefit visual verification by targeted inspection (double checking) of the diagnostic feature (shape) of the resembling (color-matching) memory object. Moreover, by attending the resembling memory object, this object may also be placed in a prioritized state, that may further benefit search by facilitating external visual guidance ^5^.

In the current study, we introduced color and shape lures to investigate the mnemonic verification process, and found that it was particularly evident following color lures. We were able to leverage this asymmetry by using the shape lures as a neutral baseline for our EEG comparisons. However, we do not wish to claim that the reported mnemonic-verification process occurs exclusively when potential targets match in color but not in shape. Ultimately, this likely depends on the precise nature of the task and stimuli. In our task, color was likely more distinguishable and therefore served as a more diagnostic feature than shape (cf. ^17,42–45^). When the external stimulus had a different color from the two memory representations (shape lure), it could easily be dismissed without prompting further inspection among the contents of working memory. However, when the external stimulus had the same color as one of the two memory objects (i.e. the color lure), it prompted additional inspection, to determine whether it *also* had the same shape. In future studies it will be interesting to parametrically vary the diagnostic values of both shape and color, as well as other potential visual features. We predict that mnemonic inspection is triggered by perceived ambiguity, regardless of whether this ambiguity is driven by shape, color, or yet other features of the target.

A direct comparison between the understudied internal verification process and the widely studied “perceptual verification” process revealed important commonalities as well as differences. We found that verification processes in both modalities were each associated with frontal-midline theta activity that was more profound when the verification process took more time to resolve. This finding aligns with the well-established role of midline frontal theta activity in cognitive control and working memory ^31–38^. At the same time, we observed notable differences. First, while mnemonic verification was predominantly reflected in microsaccades, perceptual verification was evident in larger macrosaccades, consistent with the overt inspection of one of the two visual targets on the screen. We interpret these microsaccade biases as an overt reflection (peripheral fingerprint; ^46^) of covert attention shifts within the spatial layout of working memory (as in ^23–30^). We further found a significant difference in the timing of the verification process. Inspection among the contents of working memory commenced later than inspection among the contents of perception. Thus, while perceptual and mnemonic verification processes may often be called upon together, they may not be engaged at similar speeds – at least not as reflected in our spatial gaze marker.

Despite us treating mnemonic and perceptual verification as separate, ultimately the comparison between a external visual target and internal memory representations relies on a two-way process that engages joint “mnemonic” plus “perceptual” verification. Here, we experimentally isolated the mnemonic from the perceptual verification components by selectively studying visual inspection among the contents of working memory (“mnemonic verification”), or among the content of perception (“perceptual verification”). We did this by selectively tagging either the memory or the perceptual objects-to-inspect to specific spatial locations (adopting the same logic as in ^18,19^). Yet, in real-world search, the processes of internal and external inspection in service of verification are likely often engaged concurrently and may interact in interesting, yet to be understood, ways. In future studies it will be interesting to set-up tasks to track both processes concurrently. We further point out that while our study specifically targeted search from working memory, search may also be guided by long-term memory, where external and internal inspection may also operate concurrently ^47–49^.

Our novel experimental task and analysis approach open new avenues for studying core and common components of visual search that are often indirectly inferred but rarely directly tracked. By positioning memory objects in distinct locations from the visual search object, we could leverage directional biases in microsaccades to isolate and track mnemonic inspection during visual search. Moreover, by including lures, we could track internal inspection not only when faced with actual targets, but also in response to external stimuli that were perceived as potential targets and thus called for further inspection. This approach could be expanded to study internal verification processes in more complex and elaborate search paradigms, such as hybrid search ^50,51^. This has ample potential to provide additional insights into the foundational two-way processes by which we identify what we are looking for and dismiss what we are not.

## Methods

### Participants

We performed two complementary experiments. Because the main purpose of Experiment 2 was to replicate and extend our core findings from Experiment 1, we exclusively cover Experiment-2 specific methods and results in our **Supplementary Materials**. Here, we describe the methods as they apply to Experiment 1.

Twenty-five healthy volunteers participated in the experiment (ages 18 to 46, M = 24.2, SD = 7.2; 16 females, 9 males). The sample size was determined based on previous studies from our lab with similar experimental designs and similar EEG and eye-tracking outcome variables ^23–30^. EEG data from two participants could not be used due to incomplete signal recording. The experimental procedures were reviewed and approved by the Research Ethics Committee of the Faculty of Behavioral and Movement Sciences at the Vrije Universiteit Amsterdam. All participants provided written informed consent prior to the study and were compensated with €10 per hour or equivalent credits.

### Task and Procedure

Human observers completed two versions of a visual-verification task within a single experiment: one requiring verification of whether a visual stimulus on the screen matched either of two internal visual representations held in working memory (“mnemonic verification”; Fig. 1a), and the other requiring verification of whether a single visual representation in working memory matched either of two external visual stimuli on the screen (“perceptual verification”; Fig. 3a) (akin to the approach in ^18,19^). The key distinction between the mnemonic and perceptual verification tasks lies in whether participants are comparing a single centrally presented visual target among two mnemonic representations held in working memory (enabling us to use spatial markers to isolate and track internal inspection of the left/right memory object), or comparing a single centrally encoded mnemonic target against two visual stimuli presented on the screen (enabling us to use spatial markers to isolate and track external inspection of the left/right perceptual object).

In the mnemonic-verification task, each trial (Fig. 1a) began with a memory array displaying two objects (each with distinct colors and shapes) on the left and right sides of the screen for 200 ms. This was followed by a 1000 ms delay period during which only the central fixation cross remained on the screen. Next, a memory test was presented at the center of the screen. Participants were instructed to respond whether the test object was identical (same color *and* shape) to one of the memorized objects (“same”) or different from both (“different”). The test object was chosen from the following three types with equal probability: a “target” that matched one of the memorized objects, a “color lure” that matched the color of one of the memorized objects but had a different shape, or a “shape lure” that matched the shape of one of the memorized objects but had a different color. Of the two memory objects, there was always one object that would either be a full match or a lure that matched only in color or only in shape. The other memory object was always chosen such that it did *not* match in shape *nor* in color, thus serving as the neutral object. The logic was as follows: if mnemonic verification engages visual inspection of the potential memory match (be it a target or lure), we could track this in spatial attentional deployment to the memorized location of the target or lure memory object. A “same” response was only correct for the “target” test. In trials with only a lure, a “different” response was the correct answer. The location of the target/lure and neutral memory object were counterbalanced across trials.

The trial procedure for the complementary perceptual-verification version of our task closely mirrors that of the mnemonic-verification version, with the main difference being the reversal of the memory and test array sequence (Fig. 3a). In the perceptual-verification task, participants first viewed a single memory object (search template) presented centrally. They then responded to whether this memorized object was present or absent among two test objects displayed on the left and right sides of the screen that would again contain either a target or lure on one side, and a neutral object on the other side.

In both tasks, participants pressed either the “F” or “J” key on the keyboard for the “same” and “different” (or “present” and “absent”) responses, with the key-response association counterbalanced across participants. Feedback was provided after each response, with “1” indicating a correct answer and “x” indicating an incorrect answer.

Each participant completed four sessions, with each session comprising 432 trials – yielding a total of 1728 trials per participant. Two out of four sessions involved the mnemonic-verification version of the task, while the other two sessions involved the perceptual-verification version of the task. The order of tasks was pseudo-randomized across participants. In both tasks, one of three trials contained a target, another third of trials contained a color lure, and the final third of trials contained a shape lure. This resulted in 288 trials per condition, with a total of six conditions (2 verification tasks x 3 test types). Prior to the first session, participants practiced 18 trials covering all possible conditions. The entire experiment lasted approximately 2 hours.

### Apparatus and stimuli

Stimuli were presented using MATLAB (R2022a; MathWorks) and the Psychophysics Toolbox (version 3.0.16, ^52^) on a 23-inch LED monitor with a resolution of 1920 by 1080 pixels, operating at a 240 Hz refresh rate. Participants were seated 70 cm from the screen, with their heads supported by a chin rest to ensure stable eye-tracking.

A gray background (RGB = 192, 192, 192) was used throughout the experiment, with a central fixation cross (0.75° in width and height) displayed at the center of the screen. Both memory and test objects consisted of colored shapes, randomly selected from three shapes of equal area (square [1.2° side length], circle [1.36° diameter], or diamond [1.2° side length]) and three colors (blue [RGB: 102, 255, 255], orange [RGB: 255, 153, 51], or pink [RGB: 255, 102, 255]). The two memory objects (in the mnemonic-verification task) or visual objects (in the perceptual-verification task) were displayed 5.4° to the left and right of central fixation, each with a unique shape and color.

### Analysis of behavioral data

Response accuracy was calculated as the proportion of correct responses. Reaction time (RT) was measured as the interval from the onset of the memory test to the completion of the response. Trials with RTs exceeding 2 seconds were excluded from subsequent analysis, retaining 99.44% (SD = 0.7%) of trials for further analysis.

### Eye-tracking acquisition and pre-processing

The eye tracker (EyeLink 1000, SR Research) was positioned approximately 5 cm in front of the monitor, on a table about 65 cm away from the participants’ eyes. Horizontal and vertical gaze positions were recorded for a single eye at a sampling rate of 1000 Hz. Prior to recording, the eye tracker was calibrated using the built-in calibration and validation protocols of the EyeLink software. Participants were instructed to rest their chin on the chin rest for the entire duration of each experimental block following calibration.

Offline, eye-tracking data were converted from the .edf format to .asc format and imported into MATLAB using FieldTrip^53^. Blinks were identified by detecting clusters of zeros in the eye-tracking data. Data from 100 ms before to 100 ms after each detected blink cluster were marked as Not-a-Number (NaN) to exclude residual blink artifacts. Following blink correction, the data were epoched from -200 ms to 1000 ms relative to the onset of the memory test.

### Saccade detection

For saccade detection, we utilized a velocity-based approach (cf. ^54^) that followed the procedure described in ^27^ and that we successfully employed in several prior working-memory studies from our lab (see: ^24–27,55^). Given that the memory objects in the mnemonic-verification task and the test objects in the perceptual-verification task were always presented horizontally (i.e., center-left and center-right), our saccade detection focused specifically on the horizontal channel of the eye-tracking data (as in ^24–27)^.

We first computed gaze velocity by measuring the distance between consecutive gaze positions over time. To improve precision and reduce noise, we applied temporal smoothing to the velocity data using a Gaussian-weighted moving average filter with a 7-ms sliding window, implemented via MATLAB’s “smoothdata” function. Saccade onset was identified as the first instance where the velocity exceeded a trial-specific threshold, set at 5 times the median velocity. To avoid multiple detections of the same saccade, we imposed a minimum interval of 100 ms between successive saccades.

Saccade magnitude and direction were assessed by comparing gaze positions before the saccade (from -50 to 0 ms before threshold crossing) with positions after the saccade (from 50 to 100 ms after threshold crossing). Each saccade was categorized as “toward” or “away” based on its direction (left/right) and its relation to the location of the memorized object (in the mnemonic-verification task) or the search object (in the perceptual-verification task). A “toward” saccade was aligned with the location of the target or lure-matching object while an “away” saccade was directed in the opposite direction (toward the neutral object).

After identifying and classifying the saccades, we quantified saccade rates (in Hz) using a sliding time window of 50 ms, advances in steps of 1 ms. For the primary analysis, we focused on the spatial bias in saccades, calculated as the difference in rates (in Hz) between toward and away saccades. Finally, to characterize the nature of the saccades contributing to our reported spatial modulations, we iteratively performed the above analysis as a function of saccade size (as in ^24–27,55^), considering saccade size bins of 0.5 degrees wide, sampling successive bins in -steps of 0.1 degrees.

### Saccade bias as a function of verification time

To determine if the observed saccade bias is more pronounced in trials where additional verification is required, we split trials into “fast-verification” and “slow-verification” trials using a median split of reaction times (RT). We did this separately for each condition, using condition-specific medians. This enabled us to compare the saccade bias (toward minus away) between fast and slow RT trials, separately for the different test types (target, color lure, shape lure) and tasks (mnemonic verification, perceptual verification).

### EEG acquisition and pre-processing

EEG signals were recorded using a 64-channel Biosemi system with a sampling rate of 1024 Hz. Active electrodes were positioned according to the international 10-20 system. The CMS and DRL electrodes, located on the left and right sides of POz respectively, served as the online reference. Offline, the signals were re-referenced to the average of both mastoids.

To monitor and correct for eye-movement and blink artifacts, we placed two external electrodes horizontally adjacent to the left and right eyes, and two electrodes above and below the right eye. These electrooculogram (EOG) measurements were utilized solely for data cleaning through independent component analysis (ICA), as described below. Additionally, we used an EyeLink system to collect eye-tracking data, which was used to extract our primary eye-tracking outcome measures.

Offline data analyses were performed in MATLAB using a combination of FieldTrip ^53^ and custom code. After re-referencing, the continuous EEG data were segmented into epochs from -200 to +600 ms relative to the onset of the memory test (as in ^18,19,56^). We then applied fast independent component analysis (ICA) using FieldTrip to the (concatenated) EEG epochs to identify and remove components related to blinks and eye movements. ICA components associated with eye blinks and horizontal eye movements were excluded based on their correlations with vertical electrooculography (VEOG) and horizontal electrooculography (HEOG) signals, respectively. Additionally, we used the “ft_rejectvisual.m” function in FieldTrip with the ’summary’ method to visually inspect and remove trials with exceptionally high variance, which were considered artifact trials. We did this without knowledge of the condition to which individual trials belonged. After trial removal, we retained 96.9% (SD = 1.97%) of the trials for further EEG data analysis.

### EEG time-frequency analysis

For time-frequency decomposition of the EEG data, we first applied a surface Laplacian transform to enhance the spatial resolution^57–59^. Following this, we decomposed the cleaned EEG epochs into time-frequency representations using a short-time Fourier transform with Hanning-tapered data, as implemented in FieldTrip with the ‘ft_freqanalysis.m’ function. A 300-ms sliding time window was used to estimate spectral power in the frequency range of 2 to 40 Hz, with frequency steps of 1 Hz and time steps of 10 ms.

For the analysis of our time-frequency data, we focused on activity over frontal-central electrodes where ample prior studies have reported frontal theta activity that had been implicated in working memory and cognitive control ^31–38^. Inspired by these prior findings, we focused our analysis on time-frequency activity extracted from electrodes: FCz, FC1, FC2.

Prompted by our behavioral and gaze data – that indicated robust visual inspection-for-verification for actual targets and color lures but not for shape lures – we leveraged our shape-lure condition as a neutral reference condition for our EEG comparisons. This enabled us to contrast the target and color lure conditions against this condition. We then computed the relative percentage change in time-frequency power in the following ways. For comparisons between target tests and shape lures: ((target tests – shape lures) / (target tests + shape lures)) × 100. For comparisons between color lures and shape lures: ((color lures – shape lures) / (color lures + shape lures)) × 100. In addition, To evaluate whether theta activity is more prominent in trials where additional verification is required, we additionally performed a median split on reaction times (RT) within each condition to separate fast and slow verification trials. We then calculated the percentage change in time-frequency power between slow and fast RT trials using the formula: ((slow RT – fast RT) / (slow RT + fast RT)) × 100.

Finally, to visualize topographical maps of the relative change in time-frequency power, we averaged data over the critical theta frequency band between 3-7 Hz and the time window of 200 to 600 ms (for the comparison across test types), or 400 to 600 ms (for the comparison between fast and slow verification trials) for all electrodes. Topographies merely served as an additional visualization to verify the frontal nature of the observed activity differences and were not subjected to further statistical testing.

### Statistical analysis

#### Behavioral performance data

For statistical analysis of the behavioral performance data, we used repeated-measures ANOVAs to assess response accuracy and reaction times (RTs) across the three test types (target test, color lures, and shape lures). We did this separately for accuracy and for RT. Paired sample t-tests were performed for post-hoc comparisons between each pair of test type conditions within each verification task. The Bonferroni correction method was used to adjust for multiple comparisons, and all reported p-values are Bonferroni corrected.

#### Saccade bias

To statistically analyze the temporal profiles of the spatial modulations in saccade rates (between toward and away saccades) following the onset of the test display, we utilized a cluster-based permutation approach ^60^ with the ‘ft_timelockstatistics’ function in Fieldtrip, employing the Monte Carlo method. This approach is effective for assessing data consistency across multiple adjacent time points while controlling for (i.e., bypassing) multiple comparisons. We performed 10,000 permutations to generate a single permutation distribution of the largest cluster that could occur by chance, to compare the observed clusters in the non-permuted data against. We identified clusters using Fieldtrip’s default settings, which group temporally contiguous data points showing statistical significance (*p* < 0.05) in a mass-univariate t-test. Cluster size was defined as the sum of t-values within each cluster. This method was used to evaluate the time-series data of spatial saccade bias against zero and to compare spatial saccade bias between different test types and reaction time categories (slow vs. fast RT trials).

In addition, to compare the latency of the observed spatial saccade biases between the mnemonic-and the perceptual-verification versions of the task, we normalized the saccade bias for each version of the task by dividing the observed saccade bias at each time point by the peak value of the saccade bias within that task. We then used a simplified jackknife method (as described in ^61^) to compare the onset latency of the normalized saccade bias. Onset latency was defined as the time point at which the amplitude of saccade bias reached 50% of the peak value. Paired-samples t-tests were conducted to assess differences in latency between the two versions of the task.

#### Frontal-midline theta

Statistical analysis of the EEG time-frequency employed a similar cluster-based permutation approach as in saccade-bias analyses, with this time clustering being applied in two dimensions: time *and* frequency. This method was used to compare the relative percentage change of time-frequency power across different test types (e.g., target vs. shape lure; color lure vs. shape lure) and between trials with different reaction times (e.g., slow vs. fast RT trials).

## Data and code availability

All data and analysis scripts are publicly available in the OSF: https://osf.io/4yacu/.

## Acknowledgements

This work was supported by an NWO Vidi Grant from the Dutch Research Council (14721) and an ERC Starting Grant from the European Research Council (MEMTICIPATION, 850636) to F.v.E. We thank Baiwei Liu and Anna van Harmelen for their useful discussions.

## Supplementary Materials

**Figure S1.**
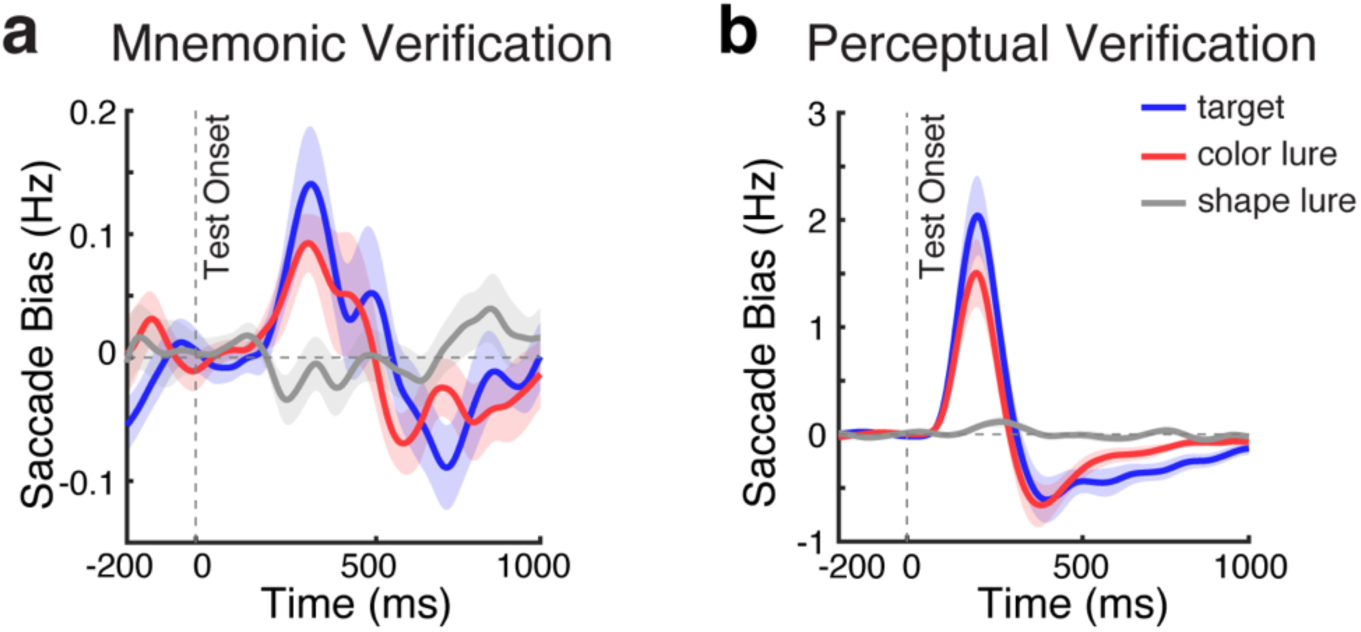
Spatial saccade bias tracks mnemonic and perceptual inspection following potential visual matches. **a)** Overlay of spatial saccade bias (toward minus away, in Hz) in the mnemonic verification task, across the three test conditions. Blue, red, and gray lines represent the gaze bias in response to actual targets, color lures, and shape lures. **b)** Overlay of spatial saccade bias in the perceptual verification across test conditions. Shadings represent ± 1 SEM across participants (n = 25).

### Supplementary Methods and Results associated with Experiment 2

The primary objective of Experiment 2 was to replicate and extend the key mnemonic-verification findings from Experiment 1. The procedure in Experiment 2 mirrors that of Experiment 1 with three notable differences. First, Experiment 2 extended Experiment 1 by introducing two additional test types in addition to “targets”, “color lures”, and “shape lures”: “color-shape lure” (in which the test stimulus matched the color of one but the shape of the other memory object; see **Fig. S2a**) and “new object”, a neutral reference condition in which the shape and the color of the test stimulus differed from both memory templates (**Fig. S2a**). Second, to ensure sufficient sensitivity with the addition of these two conditions, Experiment 2 exclusively employed the mnemonic version of our visual verification task. Finally, in Experiment 2 we did not collect EEG data.

### Methods

Although we aimed for a sample size of twenty-five participants in Experiment 2, we ended up oversampling to twenty-eight participants (ages 18 to 26, M = 21.6, SD = 2.3; 19 females, 9 males). This was not deliberate but due to a sudden urge in sign ups to the slots we had put available. The procedure followed the same mnemonic-verification task as described for Experiment 1 in the main article, though with the two additional test-types mentioned above. Each test condition was presented equally often, resulting in 216 trials per test-type. All calculations for behavioral and saccadic indices, as well as statistical analyses, followed the same methodology as in Experiment 1.

### Results

#### Behavioral results

Our behavioral analysis on response accuracy and reaction times across the five test types in Experiment 2 replicated and extended findings from Experiment 1 (**Fig. S2b**), and showed that the new objects were the easiest to dismiss, followed by shape lures, while color-shape lures were as difficult as color lures to dismiss. Repeated measures ANOVAs confirmed a significant main effect of test types (target vs. color lure vs. shape lure vs. color-shape-shift lure vs. new object) for both accuracy (*F*(4, 108) = 61.075, *p* < 0.001, η² = 0.693) and reaction time (RT) (*F*(4, 108) = 98.247, *p* < 0.001, η² = 0.784). For a full overview of all post-hoc paired-sample t-tests across different test types, see **Table S1**.

#### Saccade results

Consistent with the findings from Experiment 1, a significant saccade bias was observed following test onset for potential targets resembling the actual target (following both color-shape lures and color lures; permutation P = 0.018 for both, see **Fig. S2c,d**). Note how the saccade bias following color-shape lures is expressed relative to the location of the color-matching memory object (which was always at the other side as the shape-matching memory object in these trials). While a highly similar saccade bias was also apparant in the actual target tests, (like in Experiment 1), in Experiment 2 it did not reach statistical significance here (P = 0.143), possibly reflecting a low signal-to-noise ratio caused by an insufficient number of trials (216 trial in Experiment 2 compared to 288 trials in Experiment 1) and stronger confusion caused by more test lure types in Experiment 2 (five test types in Experiment 2 vs. three types in Experiment 1). Like in Experiment 1, the spatial saccade bias analysis as a function of saccade size showed by the saccade bias during mnemonic inspection was driven predominantly by saccades in the microsaccade range (**Fig. S2d**).

To link our saccade bias index of mnemonic inspection with verification behavior, we again split trials based on reaction times (verification times) within each condition. Similar to the findings from Experiment 1, a larger saccade bias was observed in trials requiring long vs. short verification times, though in Experiment 2 this only reached significance when testing for clusters in the actual-target condition(**Fig. S2f** permutation P < 0.001). A trend toward a more pronounced saccade bias in slow-verification trials was also seen in the color-shape-lure and color-lure conditions, although it did not reach statistical significance, possibly owing to insufficient trials.

Altogether, Experiment 2 replicated the central patterns reported in Experiment 1, such as evidence for internal inspection following potential targets like color lures and color-shape lures. At the same time, patterns in Experiment 2 were somewhat less robust than in Experiment 1. This is likely owing to the increase in the number of conditions (5 instead of 3) and, by extension, the decrease in number of trials per condition. This may have not only reduced sensitivity per condition, but possibly also resulted in more variability in the participants’ mindsets when doing the task in Experiment 2.

---

**Figure S2.**
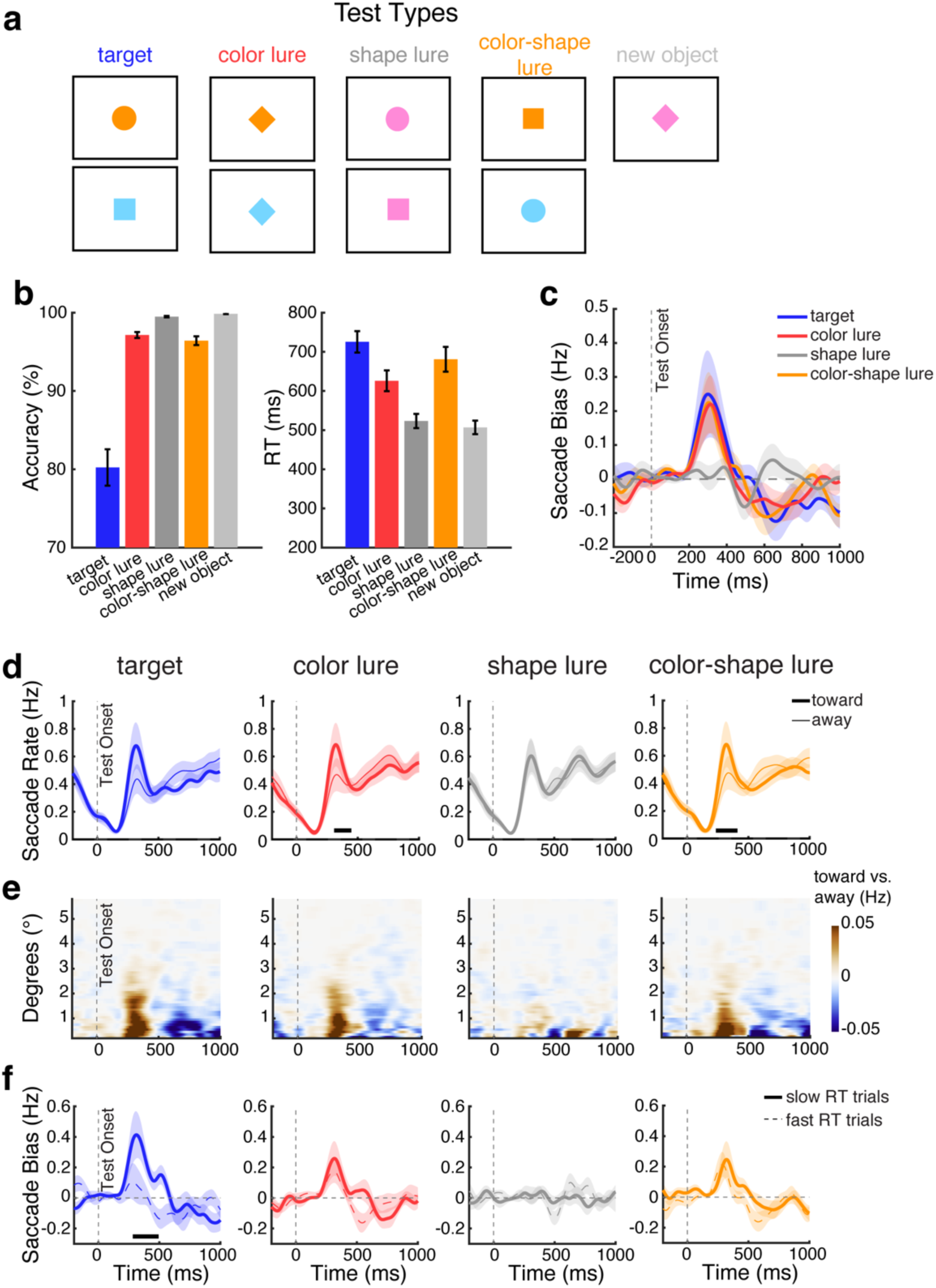
Experiment 2 replicates and expands the existence of mnemonic inspection in service of visual verification for potential visual targets. **a)** Schematic explanation of all possible test stimulus types in Experiment 2. **b)** Response accuracy (left panel) and reaction time (right panel) across test conditions. c) Spatial saccade bias (toward minus away, in Hz) across test conditions. **d)** Saccade rates (in Hz) separated by toward and away saccades relative to the location of the test stimulus that matched the memorized object, when one of the test stimuli was the actual target, color lure, shape lure, or color-shape lure (from the left to right panels). For color-shape lures, toward and away were defined relative to the color-matching object. **e)** Spatial saccade bias (toward minus away) as a function of saccade size. As in Experiment 1, the spatial saccade biases were predominantly driven by saccades in the microsacccade range (< 1 visual degree). **f)** Spatial saccade bias (toward minus away) in trials with slow RTs (solid lines) and fast RTs (dashed lines) across test conditions. Thick black horizontal lines indicate significant diTerence clusters. Shadings represent ±1 SEM across participants (n = 28).

**Table S1.** Paired-wise comparisons across di6erent test types in Experiment 2.

| Experiment 2 Test Type Effect |  |  |  |  |  |  |  |
| --- | --- | --- | --- | --- | --- | --- | --- |
| Response Accuracy |  |  |  | Reaction Time |  |  |  |
| | | $t_{(49)}$ | $p$ | | | $t_{(49)}$ | $p$ |
| Actual Target | Color Lure | -7.622** | .000 | Actual Target | Color Lure | 8.590** | .000 |
|  | Shape Lure | -8.393** | .000 |  | Shape Lure | 13.018** | .000 |
|  | Color-shape Lure | -7.193** | .000 |  | Color-shape Lure | 3.331* | .025 |
|  | New Object | -8.420** | .000 |  | New Object | 13.441** | .000 |
| Color Lure | Shape Lure | -6.062** | .000 | Color Lure | Shape Lure | 8.999** | .000 |
|  | Color-shape Lure | 1.307 | 1.000 |  | Color-shape Lure | -6.171** | .000 |
|  | New Object | -6.971** | .000 |  | New Object | 9.342** | .000 |
| Shape Lure | Color-shape Lure | 5.666** | .000 | Shape Lure | Color-shape Lure | -9.085** | .000 |
|  | New Object | -3.664* | .011 |  | New Object | 5.181** | .000 |
| Color-shape Lure | New Object | -6.128** | .000 | Color-shape Lure | New Object | 9.079** | .000 |

\*, \*\* represent significance level $p < 0.05$ and $p < 0.001$ after Bonferroni correction.

